# Extracellular appendages govern spatial dynamics and growth of *Caulobacter crescentus* on a prevalent biopolymer

**DOI:** 10.1101/2022.06.13.495907

**Authors:** Vanessa R Povolo, Glen G D’Souza, Andreas Kaczmarczyk, Astrid KM Stubbusch, Urs Jenal, Martin Ackermann

## Abstract

Microbial breakdown of carbon polymers is an essential process in all ecosystems. Carbon polymers generally require extracellular breakdown by secreted exoenzymes. Exoenzymes and breakdown products can be lost through diffusion or flow. This diffusional loss is reduced when bacteria grow in surface-associated populations where they benefit from each other’s metabolic activities. The aquatic organism *Caulobacter crescentus* was recently shown to form clonal microcolonies on the carbon polymer xylan, but to grow solitary on the monosaccharide xylose. The underlying mechanisms of this substrate-mediated microcolony formation are unknown. In particular, the importance of extracellular appendages such as pili, adhesive holdfast, and flagellum in governing the spatial arrangement of surface-grown cells is unclear. Using microfluidics coupled to automated time-lapse microscopy and quantitative image analysis, we compared the temporal and spatial dynamics of *C. crescentus* wildtype and mutant strains grown on xylan, xylose, or glucose. We found that mutants lacking type IV pili or holdfast showed altered spatial patterns in microcolonies and were unable to maintain cell densities above a threshold required for maximal growth rates on the xylan polymer, whereas mutants lacking flagella showed increased cell densities that potentially lead to increased local competition. Our results demonstrate that extracellular appendages allow bacteria to reach local cell densities that maximize single-cell growth rates in response to their nutrient environment.

## INTRODUCTION

The majority of the carbon available to bacteria in their natural environments is in the form of carbon polymers that are too large to be taken up directly by bacterial cells (*1, 2*). Bacteria secrete extracellular enzymes, such as chitinases or xylanases, to degrade polymers like chitin or xylan into oligomers and monomers that can be taken up and metabolized (*1, 2*). Microbial breakdown of carbon polymers lies at the heart of remineralization, the degradation of organic matter (*3*), and is a relevant process in the human gut (*4, 5*) and most of Earth’s ecosystems (*1*).

Secreted compounds for nutrient assimilation, such as metal chelators or exoenzymes for the breakdown of carbon polymers, as well as their breakdown products can be lost through diffusion and flow (*6*). This diffusional loss is reduced when cells grow in dense groups. In such groups, a larger fraction of the breakdown products can be taken up by cells instead of being lost by diffusion or flow (*7–10*). On the other hand, cells in spatial proximity might suffer from competition for resources (*7, 11–13*). A recent mathematical model investigated the cost-benefit ratio for secreting extracellular compounds for the acquisition of resources from the environment (*7*). According to the model, there is an intermediate cell-to-cell distance that maximizes the benefits of secretion. While competition prevails if cells are too densely packed, the synergistic effect of resource sharing declines with increasing cell-to-cell distance (*7*). In analogy, it is conceivable that to optimize growth on carbon polymers, bacteria need to maintain adequate intermediate cell densities when grown on surfaces.

If intermediate cell-to-cell distance maximizes growth rates on extracellular digested resources, one would expect bacteria to aggregate when growing on polymers. We previously tested this hypothesis with *C. crescentus*, a bacterium that is involved in the degradation of carbon polymers in a range of aquatic and terrestrial environments (*14*) and is therefore well-suited as an experimental system to investigate the role of aggregation during growth on polymers. These experiments showed that *C. crescentus* switches between aggregation and dispersal depending on whether it grows on polysaccharide or monosaccharide substrates (*10*). Observations at the single-cell level revealed that cells form clonal microcolonies on the plant polysaccharide xylan, but exhibit a planktonic lifestyle when grown on the monosaccharide xylose. Xylan is degraded by xylanases that localize to the bacterial cell envelope, and *C. crescentus* in microcolonies potentially benefit from collective xylanase activity and access to breakdown products (*10*).

Microbial aggregation is typically mediated by adhesive structures on the bacterial surface, such as pili, flagella or exopolysaccharides (*15, 16*). In *C. crescentus*, the formation of such adhesive structures is tightly coupled with its cell cycle (*17*). Each division gives rise to two morphologically and physiologically distinct cell types, a sessile stalked and a motile swarmer cell **(Fig. 1a)**. Stalked cells attach to solid substrates via an adhesive holdfast located at the tip of their stalk (*18, 19*), a polar cell extension thought to contribute to nutrient uptake (*20, 21*). Motile swarmer cells carry a flagellum and polar pili and either disperse into new environments or attach close to their stalked mothers (*17, 22*). Irreversible surface attachment of swarmer cells requires surface sensing though an active flagellum and retracting pili, culminating in the rapid synthesis of an adhesive holdfast, which anchors cells to the solid substrate (*23–27*). Thus, flagellum, pili and holdfast together coordinate *C. crescentus* surface colonization, leading to the formation of microcolonies and biofilms (*28*). The observation that holdfast production also responds to nutritional signals (*29*) argues that *C. crescentus* integrates internal and external cues to regulate surface attachment and optimize growth conditions.

**Figure 1.**
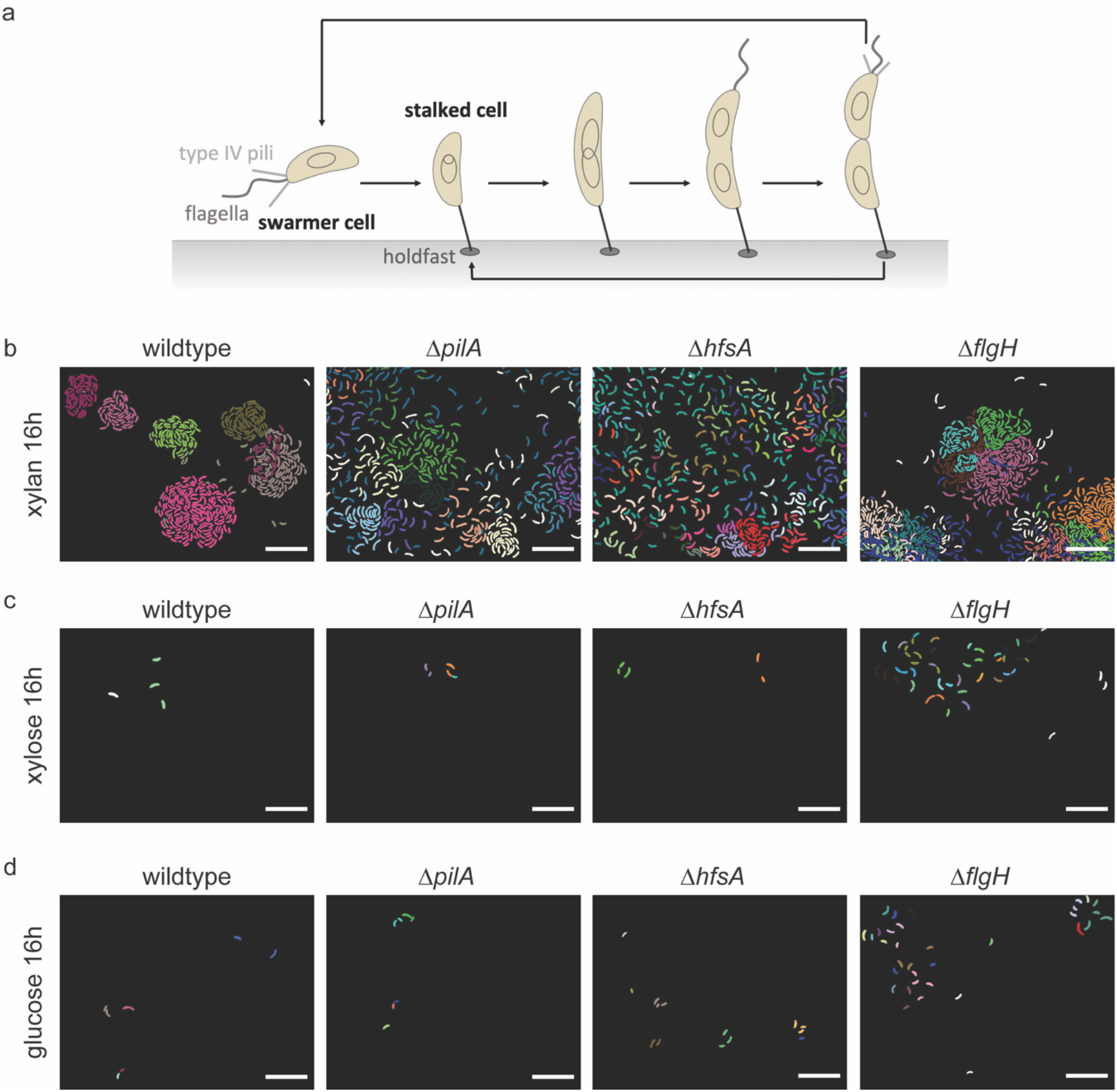
*C. crescentus* spatial growth dynamics differ in microfluidic growth chambers depending on the carbon source present and are modulated by extracellular appendages. (**a**) Schematic representation of the *C. crescentus* life cycle. Representative segmented and tracked microscopy images of *C. crescentus* CB15 wildtype and mutant strains growing on the carbon sources (**b**) xylan, (**c**) xylose, and (**d**) glucose for 16 hours in microfluidic growth chambers. Cells depicted in the same color within one image originate from the same progenitor cell. Scale bars correspond to 10 μm. The corresponding time-lapse segmented images are provided in the supplementary information **(Supplementary Movie S1-12)**.

Here, we investigate the formation of *C. crescentus* microcolonies on xylan with the specific goal to scrutinize the role of its extracellular appendages to reach optimal cell densities on surfaces for effective polymer degradation. Combining microfluidics with automated time-lapse microscopy and quantitative image analysis, we provide new insights into how surface appendages of individual microbial cells can drive the spatial organization of communities, thereby optimizing their collective growth and metabolism.

## RESULTS AND DISCUSSION

### *C. crescentus* microcolony formation on xylan depends on holdfast and type IV pili

We have recently found that the type of carbon source present influences the growth behavior of *C. crescentus* CB15 in microfluidic growth chambers (*10*). On the polymeric carbon source xylan, wildtype cells grew in dense groups of cells, so-called microcolonies, whereby all cells within a microcolony originated from a single founder cell **(Fig. 1b, wildtype)**. In contrast, on the monosaccharides xylose and glucose, cells dispersed **(Fig. 1c,d, wildtype)**. The spatial proximity of cells in microcolonies might provide a potential benefit for growth on a polymer like xylan by increasing their collective degradative activity and their access to breakdown products from neighboring cells.

Establishment of spatial proximity and the formation of microcolonies likely requires adhesive structures. To examine the role of *C. crescentus* extracellular appendages in the growth behaviors observed on different carbon sources, we used single gene knockout strains in *pilA, hfsA* or *flgH* to disrupt formation of type IV pili, holdfast or the flagellum, respectively. Type IV pili and the flagellum have been implicated in sensing initial surface contact and triggering a response that ultimately leads to irreversible attachment via c-di-GMP mediated-holdfast production (*23–27*). We performed experiments in a previously described microfluidics setup, in which growth chambers are connected to a main channel (*10, 30, 31*). Within the chambers, cells could only grow in a monolayer and were supplied with minimal salt medium containing either xylan, xylose, or glucose. We used time-lapse microscopy and automated image analysis to track the location and growth rate of the individual cells in this controlled and spatially structured microenvironment (*10, 30, 31*). In terms of natural habitats of *C. crescentus* the experimental setup resembles aquatic environments without strong flow or wet soil ecosystems (*14, 32*).

We found that extracellular appendages, such as pili and holdfast, were essential for the microcolony formation on xylan. Lack of type IV pili **(Fig. 1b, Δ*pilA*)** or a holdfast **(Fig. 1b, Δ*hfsA*)** prevented the formation of dense microcolonies on xylan (quantitative analysis of the spatial distribution is shown below). Loss of flagella, on the other hand, did not impact microcolony formation **(Fig. 1b**, Δ*flgH*), with cells showing similar spatial arrangement to wildtype cells **(Fig. 1b, wildtype)**. In contrast to the behavior on the complex polymer xylan, cells growing on the monomeric carbon sources xylose and glucose did not form microcolonies. On xylose and glucose, loss of type IV pili **(Fig. 1c,d, Δ*pilA*)** or holdfast **(Fig. 1c,d, Δ*hfsA*)** did not influence the spatial arrangement of cells compared to wildtype cells in any obvious way **(Fig. 1c,d, wildtype)**. Microcolony formation was also absent in the Δ*flgH* mutant strain **(Fig. 1c,d, Δ*flgH*)** when grown on xylose or glucose, but we observed the formation of rosettes, in which multiple cells were attached to each other through their holdfasts at the tips of their stalks (*33, 34*) **(Supplementary Figure S1)**. In rosettes, the cells are organized concentrically around their holdfasts, whereas in microcolonies cells form dense aggregates without clear concentric organization and irrespective of intercellular attachment.

Comparing the rates of cell number increase per growth chamber between the different strains growing on xylan further supported the essential role of the holdfast in surface colonization. To quantitatively describe microcolony formation on xylan, we derived the temporal dynamics of cell numbers per growth chamber using an image analysis workflow for segmentation and tracking of single cells, from which we quantified the increase in cell number over time based on an exponential model **(Supplementary Table S1)**. The cell number within a microfluidic chamber is a function of growth rate, emigration and immigration rate of cells (discussed in more detail in the next paragraph). Cell numbers in the individual growth chambers doubled every 2.8 ± 0.2 h (mean ± 95%-CI) for wildtype, 2.9 ± 0.4 h (mean ± 95%-CI) for Δ*flgH*, 3.0 ± 0.3 h (mean ± 95%-CI) for Δ*pilA*, and 3.5 ± 0.4 h for Δ*hfsA* **(Fig. 2a; Supplementary Figure S2)**. We found that Δ*hfsA* cell numbers increased significantly more slowly than those of the wild type (parametric *t*-test comparing the exponential growth parameter; *p* = 0.003; **Supplementary Table S2**), whereas Δ*pilA* and Δ*flgH* cell numbers increased similar to those of wildtype (*p* = 0.381, resp. *p* = 0.93; **Supplementary Table S2**). Thus, holdfast seems the major determinant of enhanced surface colonization on xylan, whereas type IV pili play a more accessory role in spatial patterning of microcolonies.

**Figure 2.**
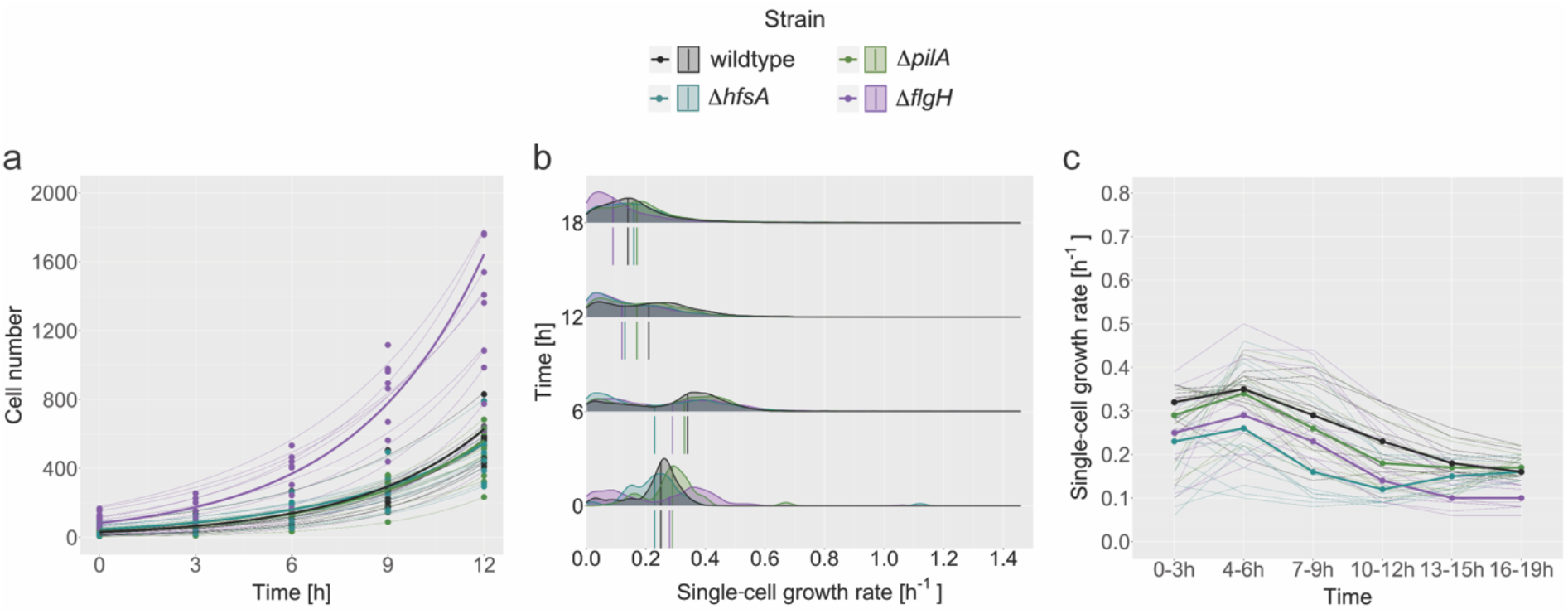
Xylan promotes surface colonization. (**a**) Cell number per microfluidic growth chamber over time for each strain on xylan. The increase in cell number for each chamber was described with an exponential model **(Supplementary Table S1)** and shown by a thin line (*R*^2^ > 0.97). Thick lines depict the exponential regression lines using the mean parameter values determined in the model functions for the individual chambers. (**b,c**) Single-cell growth rates over time for the different strains on xylan. (**b**) Density plots show the distributions of single-cell growth rates at four timepoints, i.e. 0 h, 6 h, 12 h, and 18 h for the different strains. The vertical lines within and below the density functions depict the median values for each strain. (**c**) Median values for single-cell growth rates over time for each strain on xylan **(Supplementary Table S3)**. Thin lines indicate the median single-cell growth rate trajectories of individual chambers. All graphs include data from 12 chambers, 4 per each of the 3 biological replicates, for each strain.

### Xylan promotes surface colonization

We found that the observed increase in cell number per chamber on xylan matches the expected cell number increase based on the singe-cell growth rates. Differences in the rate of increase of cell number within the chambers may arise because of differences in the cell growth rate itself, or differences in the rate of emigration when swarmer cells leave the chamber (our observations suggest that immigration is negligible, **Supplementary Movies S1-12**). In the absence of cell emigration, the doubling times derived from the observed increase in cell number within the chambers should be equal to the doubling times estimated directly from the single-cell growth rates. Emigration, on the other hand, would increase the doubling times retrieved from the observed increase in cell number within the chambers. To determine the contribution of single-cell growth rates and emigration to the overall population growth, we used the data from single-cell segmentation and tracking to estimate the single-cell growth rates based on the increase in cell area from one cell division to the next. The single-cell growth rates of all tested *C. crescentus* strains growing on xylan **(Fig. 2b,c)** reach a peak in the first six hours with values of 0.31 ± 0.04 h^-1^ (mean ± sd of all strains combined) and then decrease to values of 0.15 ± 0.03 h^-1^ (mean ± sd of all strains combined) **(Supplementary Table S3)**. Single-cell growth rates for the monosaccharides xylose and glucose are shown in the supplementary information **(Supplementary Figure S3, Supplementary Table S4 and S5).** Note that the single-cell growth rates on the monosaccharides were considerably lower than the ones on xylan. One likely reason for this is differences in the total carbon concentrations, so that growth rates in the different conditions cannot be directly compared (for details, see the Bacterial strains and growth conditions section in Materials and Methods). We derived exponential growth models from the average single-cell growth rates over the first 12 hours to obtain the predicted doubling times **(Supplementary Figure S4)** and compared these with the observed doubling times for cell numbers per chamber over the first 12 hours **(Fig. 2a; Supplementary Figure S2)**. For xylan, the doubling times predicted from the single-cell growth rates were very similar to the observed doubling times for the increase in cell number, while for xylose and glucose, observed doubling times were substantially higher than those predicted from single-cell growth rates (range of doubling time ratios across replicates: xylan, 1.08–1.15; xylose, 1.97–5.92; glucose, 2.14–3.75). An increased doubling time (corresponding to a reduced rate of cell number increase) in comparison with that predicted from the single-cell growth rates suggests higher emigration rates and hence dispersal of bacteria growing on the monosaccharides. It also suggests that xylan enhances surface colonization.

### Holdfast and type IV pili contribute to higher local cell density on xylan

The ability of *C. crescentus* cells to form dense microcolonies might contribute to their higher growth rate on xylan. To understand the temporal dynamics of microcolony formation, we generated spatial lineage trees using the position and lineage information of individual cells derived by single-cell segmentation and tracking. The spatial lineage tree for wildtype cells growing on xylan **(Fig. 3a)** showed that microcolonies were clonal and formed as a result of swarmer cells not dispersing after division. The same behavior was observed for the microcolonies formed by the mutant without a flagellum (Δ*flgH*) **(Fig. 3d)**. Loss of type IV pili (Δ*pilA*) or holdfast (Δ*hfsA*) impeded microcolony formation on xylan. The lineage trees of the Δ*pilA* **(Fig. 3b)** and Δ*hfsA* **(Fig. 3c)** mutants imply that although the swarmer cells do not settle next to their progenitor stalked cell, they tend to stay within the chamber. In this case, it is possible that the presence of xylan still serves as a cue to cells to form microcolonies, but the lack of either type IV pili or holdfast prevents swarmer cells from differentiating in proximity of the stalked cell that initiated the cell division. Type IV pili are especially important for the initial attachment of *C. crescentus* cells, while the holdfast is required for long-term stable attachment to a surface (*18, 25*). In terms of microcolony formation on xylan, lack of either type IV pili or holdfast likely prevents physical anchoring of emerging swarmer cells close to their parental stalked cells during the swarmer-to-stalked cell transition.

**Figure 3.**
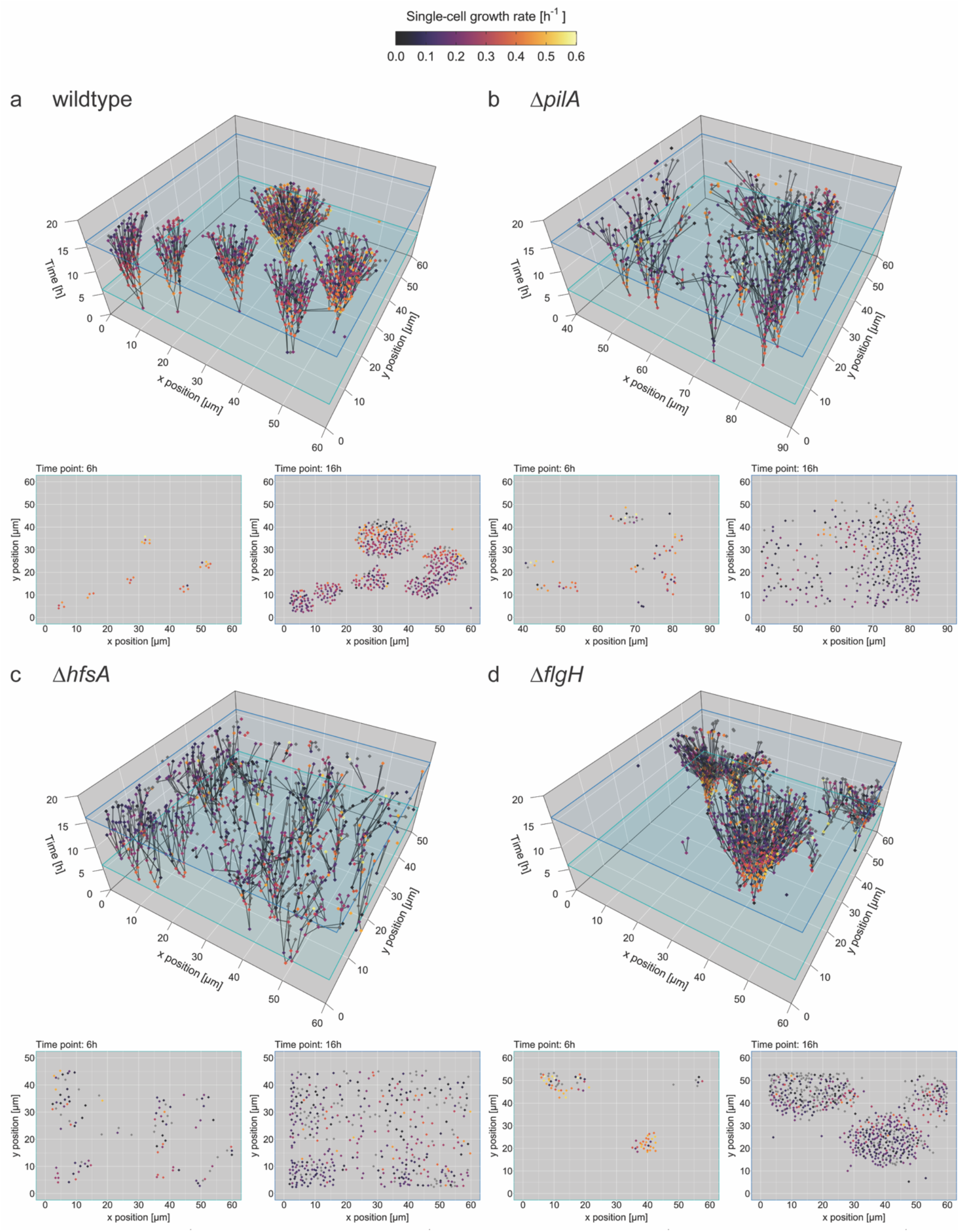
Swarmer cells stay close to their progenitor on xylan. Spatial lineage trees reconstructed for representative microfluidic growth chambers for (**a**) wildtype, (**b**) Δ*pilA*, (**c**) Δ*hfsA*, (**d**) Δ*flgH* cells growing on xylan. Single cells are plotted as dots at their specific location within the growth chamber (*x* and *y* axes) at a specific time (*z* axis) and colored according to their growth rate. Cells are connected with their progenitor cell by black lines. Branching points depict cell division events. Representative 2D planes through the spatial lineage trees at two specific time points (6 h and 16 h) are shown below the 3D plots.

These results show that the spatial distribution and hence the local cell density experienced by the individual cells is modulated by the extracellular appendages. In order to quantify the observed modulation of the spatial distribution, we derived a proxy for the spatial cell density experienced by each individual cell. To do so, we determined the number of all neighbors of each cell in the growth chamber, while giving a higher weight to close neighbors and a lower weight to more distant neighbors. We used the inverse distance squared as a weight (for details, see the Data analysis section in Materials and Methods).

The local cell density experienced by cells did indeed vary according to the carbon source and the extracellular appendages possessed by the cells. On xylan **(Fig. 4a,d)**, cells reached weighted densities of 9.23 ± 4.84 μm^-2^ (mean ± sd), while on xylose **(Fig. 4b,e)** and glucose **(Fig. 4c,f)** the weighted densities reached values of 1.2 ± 0.25 μm^-2^ and 1.01 ± 0.16 μm^-2^, respectively **(Supplementary Table S3-5)**. Weighted density differed significantly between strains on xylan (ANOVA, *F* = 13.17, *p* = 0.0048; **Supplementary Table S6**), but not on xylose (*F* = 1.64, *p* = 0.1946) or glucose (*F* = 1.22, *p* = 0.3623). The strains can be divided into two groups based on their weighted density, with wildtype and Δ*flgH* growing to significantly higher weighted density values on xylan than Δ*pilA* and Δ*hfsA (p ≤* 0.011 in all cases, Tukey’s multiple comparison test, **Supplementary Table S7**). This analysis of the weighted density supports our qualitative findings described above, that the holdfast and type IV pili are essential to form dense microcolonies on xylan. Both extracellular appendages are known to be involved in surface attachment of *C. crescentus (18, 35*), and lack of either one might prevent the swarmer cell from efficiently attaching close to the stalked cell after division.

**Figure 4.**
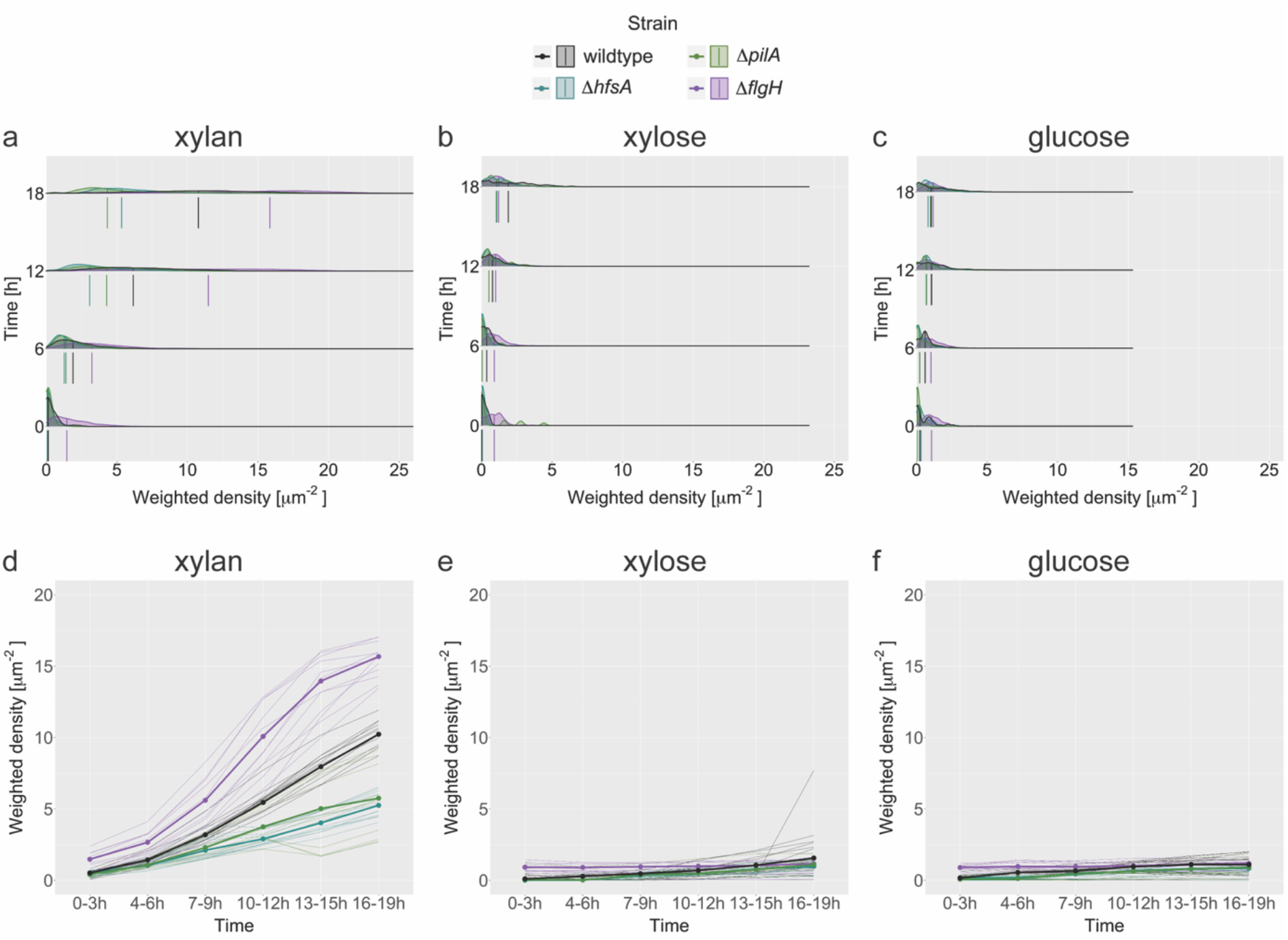
Deletion of type IV pili or holdfast decreased the cell density in microcolonies. Weighted density of single-cells over time on (**a,d**) xylan, (**b,e**) xylose, and (**c,f**) glucose for the different strains. The weighted density of a single cell was calculated by taking the sum over the inverse distance square to all other cells present. Density plots show the distributions of weighted density at four timepoints, i.e. 0 h, 6 h, 12 h, and 18 h on (**a**) xylan, (**b**) xylose, and (**c**) glucose. The vertical lines within and below the density functions depict the median values for each strain. The median weighted density over time for (**d**) xylan, (**e**) xylose, and (**f**) glucose for each strain **(Supplementary Table S3-5)**. Fine lines indicate the median weighted density of single growth chambers. All graphs include data from 12 chambers, 4 per each of the 3 biological replicates, for each strain and carbon source.

### Intermediate cell densities achieved by wildtype cells are associated with higher growth rates on xylan

Our above-described results suggested that holdfast and type IV pili influence the overall growth rate of microcolonies by retaining daughter cells close to their mother cells upon cell division. In addition, previous theoretical results (*7*) indicate that local cell density is important for the growth rate of individual cells and that intermediate densities might maximize growth rates on carbon polymers by balancing the benefits (synergy) and costs (competition) of proximity. Thus, we sought to test whether the local cell density experienced by individual cells within a microcolony also influenced their individual growth rates.

We found a link between the single-cell growth rates and the experienced local cell density of individual cells that was determined by the extracellular appendages. On xylan, single-cell growth rates reached a maximum at an intermediate weighted density of around 7.5 μm^-2^ **(Fig. 5, Supplementary Table S8)**. We found that this finding was robust to changing how the proxy for the weighted density was calculated **(Supplementary Figure S6)**. We performed the same analysis for the monosaccharides xylose and glucose and found weaker relationships between weighted density and single-cell growth rate **(Supplementary Figure S6)**.

**Figure 5.**
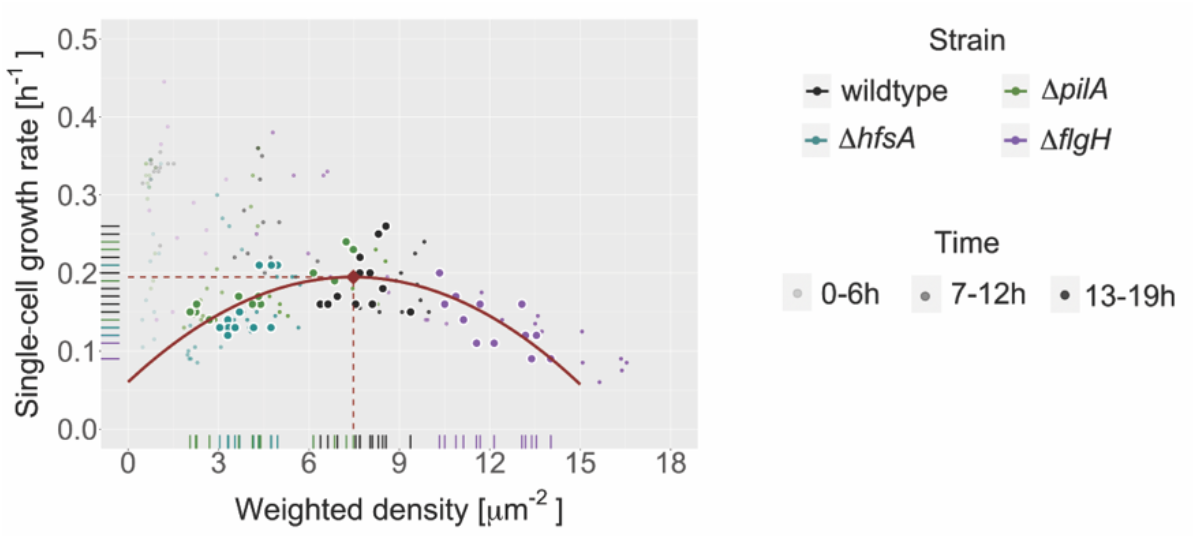
Intermediate cell densities lead to higher growth rates on xylan. Median values of single-cell growth rate and weighted density plotted against each other. Larger points with white border represent time-averages of single growth chambers. Smaller points show single growth chambers at a specific time window indicated by the degree of transparency. The distribution of the median chamber values was fitted using a quadratic regression model **(Supplementary Table S8)** and depicted in red (adj-R2 = 0.49). The found correlation is robust across different density measures **(Supplementary Figure S6)**. The maximum of the quadratic regression line is indicated with a diamond shaped point in red (y_max_ = (7.47,0.19)). The data stems from 12 chambers, 4 per each of the 3 biological replicates, for each strain.

Our results indicate that all three investigated extracellular appendages of *C. crescentus* contribute to governing the optimal cell density for the growth on the polysaccharide xylan. We found that wildtype cells reached intermediate cell densities that maximized their growth rate, while Δ*pilA* and Δ*hfsA* strains reached lower densities and hence tended to grow at lower growth rates. The Δ*flgH* mutant grew to higher densities than the wildtype and hence also tended to grow at lower growth rates **(Fig. 5)**. Differences in the single-cell growth rates between the strains were not significant (ANOVA, *p* = 0.186, **Supplementary Table S6**, Tukey’s multiple comparison test, **Supplementary Table S7**), but we found a clear trend of decreasing growth rates from wildtype over Δ*pilA* and Δ*hfsA* to Δ*flgH* **(Supplementary Figure S5)**. Our results provide evidence that type IV pili, holdfast, and flagella facilitate growth at intermediate distances that maximize growth rates of individual cells on xylan.

In conclusion, our findings imply that forming microcolonies with intermediate cell densities can be beneficial for *C. crescentus* cells, allowing them to reach maximal single-cell growth on the polysaccharide xylan. Wildtype *C. crescentus* converged around the optimal cell density, while lack of type IV pili, holdfast, or flagellum prevented cells from achieving the optimal weighted density. It is, however, important to consider that the single-cell growth rates dynamically changed over time, suggesting that additional factors than cell density and the presence or absence of the extracellular structures influence growth on xylan. Our results indicate that extracellular structures help *C. crescentus* to maintain optimal cell densities when growing on polymers in an experimental setup that recapitulates the surface-associated growth that microbes frequently experience in their natural environment (*36*).

## CONCLUSIONS

Analyzing bacterial behavior under experimental conditions mimicking important aspects of natural environments can improve our understanding of microbial ecology. In combination with genetic model systems, such an approach allows investigating the functional relevance of specific molecular, cellular and behavioral traits for the overall growth and survival of bacteria in their environment. *C. crescentus* is an extensively studied extensively model system that has resulted in the discovery of many regulatory processes involved in coordinating growth, division and behavior (*37, 38*). In contrast, the understanding of how *C. crescentus* optimizes growth and survival in its natural environment is still limited. In this study, we aimed to shed further light into changes in bacterial growth dynamics in response to different nutrient environments and identify the involved cellular structures. Our data suggests that depending on the carbon source different spatial cell densities result in maximal single-cell growth. Optimal density is likely more relevant for nutrients that require extracellular degradation, such as the studied carbon polymer xylan studied here. Furthermore, we demonstrated that extracellular appendages involved in surface adhesion impact the spatial proximity of cells. In the natural habitat of bacteria, most carbon sources require extracellular breakdown (*1, 2*) and the ability of growing in spatial proximity likely makes this degradation process more efficient. Microbial decomposition of carbon polymers is essential for the global biogeochemical cycles (*3*) and an important process in the human gut (*4, 5*). Our study provided a better understanding of how bacteria adapt to different nutrient sources, such as monosaccharides and polysaccharides, to balance benefits and costs of group formation.

## MATERIALS AND METHODS

### Bacterial strains and growth conditions

Experiments were performed using *Caulobacter crescentus* CB15 wildtype (*10*), Δ*pilA (39), DhfsA* (UJ9035 from Urs Jenal, University of Basel, Switzerland), and Δ*flgH* (UJ8848 from Urs Jenal, University of Basel, Switzerland) mutants. We used the pXGFPC-2 P*lac*::mKate2 plasmid (*22*) to introduce a fluorescent phenotypic marker by electroporation, as described previously (*22*). Chromosomal recombinants were selected using kanamycin resistance and tested for fluorescence expression.

Cells were routinely grown on Peptone Yeast Extract (PYE) agar supplemented with 20 μg/ml kanamycin or in M2 minimal salts medium containing xylose (0.05% w/v), glucose (0.05% w/v), or xylan (0.1% w/v). Stock solutions of xylose (20% w/v, Sigma Aldrich), glucose (20% w/v, Sigma Aldrich), and xylan (2% w/v, Megazyme) were prepared with nanopure water and filter sterilized using 0.4 μm surfactant-free cellulose acetate filters (Corning). Overnight cultures were prepared by inoculating a 15 ml culture tube containing M2 minimal salts medium containing glucose (M2G) with a single bacterial colony from PYE agar.

### Microfluidics

Microfluidics experiments were performed as described previously (*10, 30, 31*). Microfluidic devices were designed with chambers of 60 × 60 x 0.56 μm or 60 x 120 x 0.56 μm (length x width x height) that were open on one side (60 or 120 μm) to a 100 μm wide feeding channel of 22 μm height. Polydimethylsiloxane elastomers (PDMS, Sylgard 194 Silicone Elastomer Kit, Dow Corning) were prepared with a 1:8 ratio and poured onto a wafer. The mix was degassed using a desiccator and then baked for 2 h at 80 °C for curing. PDMS chips were cut out using a scalpel. Inlets and outlets of 0.75 mm diameter for medium were punched into the feeding channel. PDMS chips were bound to round glass coverslips of 50 mm diameter (No. 1, Menzel-Gläser) using a Plasma Cleaner (PDC-32G-2, Harrik Plasma) for 30 s at maximum power. A thermal plate at 100 °C was used to stabilize the bonding.

Overnight cultures of *C. crescentus* CB15 strains were grown at 30 °C with shaking (220 rpm) in M2G to an OD_600_ of around 0.2. Aliquots of 1 ml of this cell suspension were centrifuged (13’000 rpm, 2 min), the cells were washed twice with M2 minimal salts medium and resuspended in 100 μl of M2 minimal salts medium. Using a 10 μl pipette, cells were loaded into the PDMS chip.

To provide the carbon source during experiments, 50 ml luer lock syringes (Pic Solution) were filled with M2 minimal salts media containing either xylose (0.001% w/v), glucose (0.001% w/v), or xylan (0.1% w/v) and loaded onto single- or multi-channel syringe pumps (NE-300, NE-1600 or NE-1800, New Era Pump Systems). The PDMS chip was connected with the syringes containing the growth media using a combination of 20-G needles (0.9 × 70 mm, Huberlab), larger tubing (Tygon microbore S54HL, ID 0.76 mm, OD 2.29 mm, Fisher Scientific), and small tubing (Adtek, ID 0.3 mm, OD 0.76 mm, Fisher Scientific).

The inlets of the PDMS chip were connected to the syringes containing the growth media after loading the cells and small tubing was used to connect the outlets with a waste bottle. A flow rate of 0.1 ml/h was used for all experiments to ensure constant nutrient supply of the chambers via diffusion from the channel.

### Time-lapse microscopy

An Olympus IX83 inverted microscope system with automated stage controller (Marzhauser Wetzlar), shutters, and laser-based autofocus systems (Olympus ZDC 2) was used for microscopy imaging. Several positions on one PDMS chip were imaged in parallel. Phase-contrast and fluorescence images for every position were taken at 7 min intervals. An UPLFLN 100× oil immersion objective (Olympus) and an ORCA-flash 4.0 v2 or v4 sCMOS camera (Hamamatsu, Japan) was used for image acquisition. Fluorescence imaging was performed using a X-Cite120 120 W high pressure metal halide arc lamp (Lumen Dynamics) with a TXRED fluorescent filter (Chroma). A cellVivo microscope incubation system (Pecon GmbH, Germany) or Cube incubation system (Life Imaging Services, Switzerland) maintained the growth conditions in the course of an experiment at 30 °C.

### Image analysis

Fluorescent channel images were first converted to tiff format using a custom-scripted macro for Fiji/ImageJ v2.0 (*40*) and later aligned and cropped to the boundaries of the microfluidic growth chambers with Matlab v2019b using SuperSegger (*41*). The resulting images were deconvolved using a point-spread function (*42*) and fed into ilastik v1.3.2 (*43*) for segmentation and tracking of single cells. Further analysis of the data was performed in R Studio v2021.09.1 (*44*) with R v4.1.2. Single-cell growth rates were calculated from the increase of cell area of a single cell over time by taking the slope of the linearized fitted exponential model. The weighted cell density was calculated with the R package *spatstat* v2.2.0 (*45*) by taking for every cell the sum over the inverse distance square to all other cells in the chamber for each frame. This was motivated by the fact that cells are metabolically coupled through processes that rely on diffusion of enzymes, nutrients and metabolites. While the number and properties of the diffusing compounds are not known in detail, using the inverse distance squared takes into account the fact that the diffusional coupling between cells decreases rapidly with their distance. Alternative proxies to determine the weighted density were calculated the same way using the inverse distance or the inverse distance cubed instead of the inverse distance squared. Lineage trees were generated with the R package *plot3Drgl* v1.0.2. Visualization of ridged plots was done with the R package *ggridges* v0.5.3. For all other plots a combination of the R packages *ggplot2* v3.3.5 and *ggpubr* v0.4.0.

### Datasets and statistical analysis

Microfluidics experiments were performed in 3 biological replicates. From each of the replicates, 4 of the 8 imaged microfluidic growth chambers per strain and per carbon source were randomly chosen. In total, 12 microfluidic growth chambers for each strain on each carbon source were analyzed. For xylan, 28’723 wildtype, 20’195 Δ*pilA*, 19’586 Δhfs*A*, and 37’776 Δ*flgH* cells were analyzed; for xylose, 436 wildtype, 421 Δ*pilA*, 388 Dhfs*A*, and 1’355 Δ*flgH* cells were analyzed; and for glucose, 938 wildtype, 652 Δ*pilA*, 663 Dhfs*A*, and 2’197 Δ*flgH* cells were analyzed. For statistical analysis of the effect of extracellular appendage knockout on local cell density and single-cell growth rate, cells were aggregated at the growth chamber level by taking the median values for each growth chamber. The data was log-transformed and a linear mixed-effects model was fitted for each carbon source. The model integrated the separate experiments and growth chambers as random effects and the strains as fixed effect. Statistical analysis was performed in R Studio v2021.09.1 (*44*) with R v4.1.2 using the R packages *lmerTest* v3.1.3, *lme4* v1.1.27.1, and *multcomp* v1.4.17. In comparisons, a *p*-value < 0.05 was considered significant.

## ACKNOWLEDGMENTS

We thank Johannes Keegstra, Jonasz Słomka, and Nicola Zamboni for helpful feedback and discussions, Zemer Gitai for the fluorescent phenotypic marker plasmid. Further we thank Russell Naisbit for editorial support for the manuscript. This project was funded by the Swiss National Science Foundation (grant number 31003A_169978) to MA, an ETH fellowship and a Marie Curie Actions for People COFUND program fellowship (FEL-37-16-1) to GD, an ETH Career Seed Grant (SEED-14 18-1) to GD, the Simons Foundation Collaboration on Principles of Microbial Ecosystems (PriME grant number 542389) to MA, and by ETH Zurich and Eawag.

## Author contributions

VP conceptualized the research and designed the experiments with input from GD, MA, AK, and UJ. AK and UJ provided the mutant strains. VP performed all experiments and analyzed the data with advice from GD and MA. AS developed the single-cell growth rate computation for tracked cells. VP wrote the manuscript with input from GD, AK, AS, UJ, and MA.

