## Supplementary Information for "Extracellular appendages govern spatial dynamics and growth of *Caulobacter crescentus* on a prevalent biopolymer"

**This file includes:**

Supplementary Tables 1 to 8

Supplementary Figures 1 to 6

Supplementary Videos 1 to 12

### SUPPLEMENTARY TABLES

| CS | Strain | N | Exponential parameters | | | | pRsq | | Doubling time [h] | |
|---|---|---|---|---|---|---|---|---|---|---|
| | | | **a mean** | **a range** | **b mean** | **b range** | **mean** | **range** | **mean** | **95% CI** |
| xylan | wildtype | 12 | 31.14 | [11,86] | 0.25 | [0.19,0.31] | 0.998 | [0.9951,0.9995] | 2.812 | ± 0.24 |
| | Δ*pilA* | 12 | 33.07 | [5,54] | 0.24 | [0.19,0.32] | 0.997 | [0.9934,0.9997] | 3.013 | ± 0.32 |
| | Δ*hfsA* | 12 | 47.20 | [13,91] | 0.20 | [0.16,0.26] | 0.992 | [0.9735,0.9961] | 3.494 | ± 0.37 |
| | Δ*flgH* | 12 | 83.15 | [7,180] | 0.25 | [0.18,0.34] | 0.994 | [0.9723,0.9995] | 2.934 | ± 0.43 |
| xylose | wildtype | 12 | 6.82 | [1,22] | 0.06 | [-0.04,0.18] | 0.646 | [0.0283,0.9746] | 20.25 | ± 31.5 |
| | Δ*pilA* | 12 | 7.6 | [1,26] | 0.05 | [-0.01,0.1] | 0.531 | [0.0205,0.9035] | 14.88 | ± 17 |
| | Δ*hfsA* | 12 | 4.23 | [2,10] | 0.07 | [0.03,0.11] | 0.728 | [0.3129,0.9656] | 11.3 | ± 3.53 |
| | Δ*flgH* | 12 | 46.62 | [8,113] | 0.02 | [0.01,0.07] | 0.347 | [0.0409,0.8904] | 66.69 | ± 25.5 |
| glucose | wildtype | 12 | 16.13 | [5,28] | 0.05 | [-0.02,0.1] | 0.842 | [0.0587,0.9748] | 13.82 | ± 14.5 |
| | Δ*pilA* | 12 | 12.91 | [4,29] | 0.04 | [-0.01,0.07] | 0.764 | [0.2524,0.9508] | 12.15 | ± 15.1 |
| | Δ*hfsA* | 12 | 16.24 | [3,42] | 0.03 | [-0.07,0.09] | 0.637 | [0.01,0.936] | 28.24 | ± 34.8 |
| | Δ*flgH* | 12 | 50.53 | [22,83] | 0.03 | [0.002,0.04] | 0.787 | [0.0143,0.9856] | 52.96 | ± 69.8 |

**Table S1 | Summary statistics of exponential model for cell numbers.** Increase in cell numbers per chamber in the first 12 hours was exponential and best described by the following model: cell number ~ a*exp(b*time). CS, carbon source. N, number of chambers. pRsq, pseudo R squared calculated by 1-RSS/TSS. 95% CI, 95% confidence interval.

| CS | Strain | wildtype | Δ*pilA* | Δ*hfsA* | Δ*flgH* |
|---|---|---|---|---|---|
| xylan | wildtype | - | 0.3808 | 0.0027 | 0.9303 |
| | Δ*pilA* | 0.3808 | - | 0.0540 | 0.5703 |
| | Δ*hfsA* | 0.0027 | 0.0540 | - | 0.0364 |
| | Δ*flgH* | 0.9303 | 0.5703 | 0.0364 | - |

**Table S2 | P-values from parametric t-test for the b parameter of the exponential models for cell number over time.** CS, carbon source.

| CS | Strain | Time | N | Single-cell growth rate Median [h$^{-1}$] | Single-cell growth rate IQR [h$^{-1}$] | Single-cell growth rate CV [%] | Weighted density Median [μm$^{-2}$] | Weighted density IQR [μm$^{-2}$] | Weighted density CV [%] |
|---|---|---|---|---|---|---|---|---|---|
| xylan | wildtype | 0-3h | 12’514 | 0.32 | 0.19 | 44.74 | 0.5227 | 0.6452 | 128.76 |
| | | 4-6h | 25’777 | 0.35 | 0.24 | 46.36 | 1.4331 | 1.6518 | 93.73 |
| | | 7-9h | 59’500 | 0.29 | 0.28 | 61.72 | 3.1973 | 2.7282 | 64.48 |
| | | 10-12h | 125’960 | 0.23 | 0.24 | 72.90 | 5.4581 | 3.8821 | 51.87 |
| | | 13-15h | 232’264 | 0.18 | 0.17 | 79.19 | 7.9593 | 4.9589 | 44.40 |
| | | 16-19h | 308’346 | 0.16 | 0.13 | 82.34 | 10.2306 | 5.4223 | 37.86 |
| | Δ*pilA* | 0-3h | 12’456 | 0.29 | 0.24 | 50.51 | 0.4088 | 0.5052 | 103.14 |
| | | 4-6h | 25’078 | 0.34 | 0.3 | 52.99 | 1.0742 | 1.0192 | 81.68 |
| | | 7-9h | 56’994 | 0.26 | 0.29 | 70.58 | 2.2962 | 1.9019 | 66.32 |
| | | 10-12h | 115’946 | 0.18 | 0.22 | 87.99 | 3.7537 | 3.2056 | 59.22 |
| | | 13-15h | 170’862 | 0.17 | 0.16 | 78.93 | 5.0254 | 4.8880 | 60.78 |
| | | 16-19h | 209’425 | 0.17 | 0.14 | 85.75 | 5.7539 | 5.5498 | 94.50 |
| | Δ*hfsA* | 0-3h | 16’513 | 0.23 | 0.27 | 67.38 | 0.5017 | 0.5927 | 127.57 |
| | | 4-6h | 31’112 | 0.26 | 0.33 | 72.23 | 1.0419 | 1.0475 | 98.66 |
| | | 7-9h | 63’405 | 0.16 | 0.28 | 115.94 | 2.1118 | 2.1278 | 78.33 |
| | | 10-12h | 114’539 | 0.12 | 0.2 | 139.82 | 2.9068 | 2.6166 | 69.86 |
| | | 13-15h | 187’033 | 0.15 | 0.18 | 97.87 | 4.0240 | 2.9540 | 55.24 |
| | | 16-19h | 241’928 | 0.16 | 0.16 | 104.72 | 5.2539 | 3.2564 | 46.35 |
| | Δ*flgH* | 0-3h | 30’227 | 0.25 | 0.28 | 69.94 | 1.4783 | 1.4960 | 84.27 |
| | | 4-6h | 59’583 | 0.29 | 0.31 | 69.86 | 2.6636 | 2.4533 | 75.25 |
| | | 7-9h | 133’539 | 0.23 | 0.28 | 86.11 | 5.6092 | 4.8234 | 63.94 |
| | | 10-12h | 270’908 | 0.14 | 0.21 | 123.37 | 10.0819 | 7.1577 | 48.01 |
| | | 13-15h | 420’323 | 0.1 | 0.14 | 139.41 | 13.9565 | 7.3917 | 45.52 |
| | | 16-19h | 459’112 | 0.1 | 0.14 | 151.87 | 15.6684 | 6.6788 | 195.78 |

**Table S3 | Summary statistics weighted density and single-cell growth rates on xylan.** CS, carbon source. N, number of cells. IQR, interquartile range. CV, coefficient of variation.

| CS | Strain | Time | N | Single-cell growth rate Median [h$^{-1}$] | Single-cell growth rate IQR [h$^{-1}$] | Single-cell growth rate CV [%] | Weighted density Median [μm$^{-2}$] | Weighted density IQR [μm$^{-2}$] | Weighted density CV [%] |
|---|---|---|---|---|---|---|---|---|---|
| xylose | wildtype | 0-3h | 1’962 | 0.12 | 0.14 | 74.61 | 0.1050 | 0.5038 | 690.22 |
| | | 4-6h | 2’175 | 0.09 | 0.14 | 125.26 | 0.2840 | 0.5175 | 162.21 |
| | | 7-9h | 2’604 | 0.07 | 0.17 | 152.45 | 0.4696 | 0.6818 | 137.35 |
| | | 10-12h | 3’123 | 0.09 | 0.18 | 111.90 | 0.6923 | 1.0260 | 125.72 |
| | | 13-15h | 4’071 | 0.15 | 0.17 | 68.80 | 1.0648 | 1.7095 | 119.71 |
| | | 16-19h | 4’584 | 0.18 | 0.16 | 75.52 | 1.5638 | 2.4545 | 119.09 |
| | Δ*pilA* | 0-3h | 2’479 | 0.09 | 0.06 | 67.39 | 0.0443 | 0.2986 | 1376.85 |
| | | 4-6h | 2’204 | 0.09 | 0.06 | 56.84 | 0.0269 | 0.5029 | 1656.95 |
| | | 7-9h | 2’682 | 0.08 | 0.08 | 93.45 | 0.3933 | 0.5994 | 133.32 |
| | | 10-12h | 3’392 | 0.08 | 0.08 | 108.79 | 0.4950 | 0.8280 | 151.71 |
| | | 13-15h | 4’425 | 0.09 | 0.08 | 72.44 | 0.7644 | 1.1750 | 110.78 |
| | | 16-19h | 4’503 | 0.09 | 0.08 | 89.16 | 1.1021 | 1.2719 | 85.03 |
| | Δ*hfsA* | 0-3h | 1’523 | 0.11 | 0.06 | 44.47 | 0.0114 | 0.0541 | 2257.22 |
| | | 4-6h | 1’607 | 0.11 | 0.07 | 46.57 | 0.0235 | 0.1725 | 1516.00 |
| | | 7-9h | 2’195 | 0.13 | 0.09 | 49.28 | 0.3444 | 0.4566 | 100.14 |
| | | 10-12h | 2’941 | 0.16 | 0.11 | 42.31 | 0.4838 | 0.5964 | 116.79 |
| | | 13-15h | 3’971 | 0.16 | 0.1 | 43.69 | 0.7944 | 0.8918 | 91.85 |
| | | 16-19h | 4’315 | 0.16 | 0.1 | 47.41 | 0.9767 | 1.0308 | 82.63 |
| | Δ*flgH* | 0-3h | 13’244 | 0.05 | 0.06 | 119.99 | 0.9219 | 0.8759 | 90.37 |
| | | 4-6h | 14’277 | 0.05 | 0.05 | 114.00 | 0.9080 | 0.8035 | 91.29 |
| | | 7-9h | 15’574 | 0.05 | 0.05 | 123.70 | 0.9527 | 0.8246 | 91.89 |
| | | 10-12h | 16’677 | 0.06 | 0.05 | 117.01 | 0.9976 | 0.8563 | 97.11 |
| | | 13-15h | 19’178 | 0.06 | 0.09 | 135.01 | 1.0445 | 0.9238 | 104.93 |
| | | 16-19h | 18’599 | 0.07 | 0.13 | 159.73 | 1.1686 | 0.9836 | 100.67 |

**Table S4 | Summary statistics weighted density and single-cell growth rates on xylose.** CS, carbon source. N, number of cells. IQR, interquartile range. CV, coefficient of variation.

| CS | Strain | Time | N | Single-cell growth rate Median [h$^{-1}$] | IQR [h$^{-1}$] | CV [%] | Weighted density Median [μm$^{-2}$] | IQR [μm$^{-2}$] | CV [%] |
|---|---|---|---|---|---|---|---|---|---|
| glucose | wildtype | 0-3h | 4'491 | 0.15 | 0.15 | 53.56 | 0.1704 | 0.5674 | 391.58 |
| | | 4-6h | 6'428 | 0.12 | 0.13 | 78.43 | 0.5560 | 0.7458 | 146.58 |
| | | 7-9h | 7'842 | 0.11 | 0.13 | 83.56 | 0.6636 | 0.6489 | 117.05 |
| | | 10-12h | 8'958 | 0.11 | 0.12 | 77.87 | 0.9602 | 1.1057 | 104.38 |
| | | 13-15h | 10'476 | 0.11 | 0.08 | 62.90 | 1.1172 | 1.4530 | 103.11 |
| | | 16-19h | 11'239 | 0.1 | 0.08 | 87.64 | 1.1047 | 1.6821 | 113.31 |
| | Δ*pilA* | 0-3h | 3'886 | 0.09 | 0.07 | 49.42 | 0.0812 | 0.3516 | 712.65 |
| | | 4-6h | 4'530 | 0.1 | 0.08 | 55.32 | 0.1280 | 0.6866 | 580.67 |
| | | 7-9h | 5'489 | 0.09 | 0.1 | 67.39 | 0.4929 | 0.8242 | 156.11 |
| | | 10-12h | 6'464 | 0.09 | 0.09 | 72.26 | 0.6363 | 0.8226 | 135.13 |
| | | 13-15h | 7'750 | 0.1 | 0.08 | 76.12 | 0.7920 | 1.1186 | 123.73 |
| | | 16-19h | 8'540 | 0.11 | 0.08 | 61.96 | 0.9184 | 1.3489 | 115.35 |
| | Δ*hfsA* | 0-3h | 4'654 | 0.1 | 0.06 | 47.33 | 0.1824 | 0.4131 | 240.39 |
| | | 4-6h | 4'401 | 0.1 | 0.06 | 43.81 | 0.1987 | 0.5253 | 231.13 |
| | | 7-9h | 5'003 | 0.09 | 0.07 | 54.06 | 0.4413 | 0.6200 | 117.39 |
| | | 10-12h | 6'465 | 0.1 | 0.08 | 51.66 | 0.6729 | 0.6052 | 98.68 |
| | | 13-15h | 7'561 | 0.1 | 0.07 | 50.68 | 0.8253 | 0.8519 | 85.50 |
| | | 16-19h | 7'121 | 0.1 | 0.07 | 68.52 | 0.8450 | 0.8911 | 87.73 |
| | Δ*flgH* | 0-3h | 14'296 | 0.09 | 0.12 | 94.59 | 0.9000 | 0.8945 | 102.29 |
| | | 4-6h | 18'153 | 0.08 | 0.11 | 92.44 | 0.9772 | 0.9576 | 90.26 |
| | | 7-9h | 19'773 | 0.07 | 0.1 | 104.51 | 0.9617 | 0.9428 | 103.33 |
| | | 10-12h | 21'814 | 0.07 | 0.09 | 108.35 | 1.0174 | 0.9371 | 88.62 |
| | | 13-15h | 25'447 | 0.08 | 0.08 | 91.12 | 1.1030 | 0.9502 | 81.70 |
| | | 16-19h | 27'362 | 0.08 | 0.09 | 130.02 | 1.1780 | 1.0180 | 80.38 |

**Table S5 | Summary statistics weighted density and single-cell growth rates on glucose.** CS, carbon source. N, number of cells. IQR, interquartile range. CV, coefficient of variation.

| CS | Weighted density Sum Sq / Mean Sq | numDF / denDF | F value | *p*-value | Single-cell growth rate Sum Sq / Mean Sq | numDF / denDF | F value | *p*-value |
|---|---|---|---|---|---|---|---|---|
| xylan | 0.4914 / 0.1638 | 3 / 5.9997 | 13.173 | 0.004756 | 0.0507 / 0.0169 | 3 / 6 | 2.2241 | 0.186 |
| xylose | 2.9957 / 0.9986 | 3 / 42 | 1.6397 | 0.1946 | | | | |
| glucose | 2.72 / 0.9067 | 3 / 8 | 1.2244 | 0.3623 | | | | |

**Table S6 | Parametric ANOVA.** CS, carbon source. Sum Sq, sum of squares. Mean Sq, mean squares. numDF, degrees of freedom in the numerator. denDF, degrees of freedom in the denominator.

| CS | Strain | Weighted density | | | | Single-cell growth rate | | | |
|---|---|---|---|---|---|---|---|---|---|
| | | wildtype | Δ*pilA* | Δ*hfsA* | Δ*flgH* | wildtype | Δ*pilA* | Δ*hfsA* | Δ*flgH* |
| xylan | wildtype | - | 0.01097 | 0.00755 | 0.159 | - | 0.9682 | 0.561 | 0.0773 |
| | Δ*pilA* | 0.01097 | - | 0.99949 | < 0.001 | 0.9682 | - | 0.8325 | 0.2107 |
| | Δ*hfsA* | 0.00755 | 0.99949 | - | < 0.001 | 0.561 | 0.8325 | - | 0.6922 |
| | Δ*flgH* | 0.159 | < 0.001 | < 0.001 | - | 0.0773 | 0.2107 | 0.6922 | - |
| xylose | wildtype | - | 0.997 | 0.574 | 0.812 | | | | |
| | Δ*pilA* | 0.997 | - | 0.447 | 0.904 | | | | |
| | Δ*hfsA* | 0.574 | 0.447 | - | 0.132 | | | | |
| | Δ*flgH* | 0.812 | 0.904 | 0.132 | - | | | | |
| glucose | wildtype | - | 0.86 | 0.603 | 0.957 | | | | |
| | Δ*pilA* | 0.86 | - | 0.97 | 0.565 | | | | |
| | Δ*hfsA* | 0.603 | 0.97 | - | 0.3 | | | | |
| | Δ*flgH* | 0.957 | 0.565 | 0.3 | - | | | | |

**Table S7 | P-values of Tukey's multiple comparison test.** CS, carbon source.

| Density measure | CS | N | Model parameters | | | | Rsq | | vertex | |
|---|---|---|---|---|---|---|---|---|---|---|
| | | | a | *p*-value of a | b | c | mRsq | adjRsq | $y_{max}$ | $x_{ymax}$ |
| Weighted density: $\sum_{i=1}^{n}\frac{1}{d_i^2}$ | xylan | 48 | -0.002 | $8.79e^{-8}$ | 0.036 | 0.06 | 0.488 | 0.465 | 0.195 | 7.473 |
| | xylose | 48 | -0.013 | 0.754 | 0.053 | 0.074 | 0.065 | 0.024 | 0.127 | 1.611 |
| | glucose | 48 | -0.094 | 0.001 | 0.161 | 0.033 | 0.337 | 0.308 | 0.102 | 0.831 |
| $\sum_{i=1}^{n}\frac{1}{d_i}$ | xylan | 48 | $-4.49e^{-5}$ | 0.0001 | 0.004 | 0.092 | 0.38 | 0.353 | 0.183 | 45.538 |
| | xylose | 48 | -0.004 | 0.113 | 0.019 | 0.088 | 0.077 | 0.036 | 0.113 | 2.69 |
| | glucose | 48 | -0.004 | $8.41e^{-5}$ | 0.032 | 0.044 | 0.313 | 0.283 | 0.106 | 3.806 |
| $\sum_{i=1}^{n}\frac{1}{d_i^3}$ | xylan | 48 | -0.009 | $1.10e^{-5}$ | 0.061 | 0.095 | 0.352 | 0.324 | 0.195 | 3.128 |
| | xylose | 48 | -0.029 | 0.75 | 0.128 | 0.065 | 0.185 | 0.149 | 0.166 | 1.039 |
| | glucose | 48 | -0.092 | 0.321 | 0.152 | 0.049 | 0.291 | 0.259 | 0.112 | 0.747 |

**Table S8 | Summary statistics of quadratic model for correlation between single-cell growth rate and cell density measures.** Cell density was estimated using different sums over the inverse distance *d* between cells. The correlation between single-cell growth rate (y) and cell density (x) was described by the following model: y ~ a*x^2 + b*x + c. CS, carbon source. N, number of chambers. Rsq, R squared. mRsq, multiple R squared. adjRsq, adjusted R squared. $y_{max}$, maximum of quadratic formula. $x_{ymax}$, x value corresponding to $y_{max}$.

### SUPPLEMENTARY FIGURES

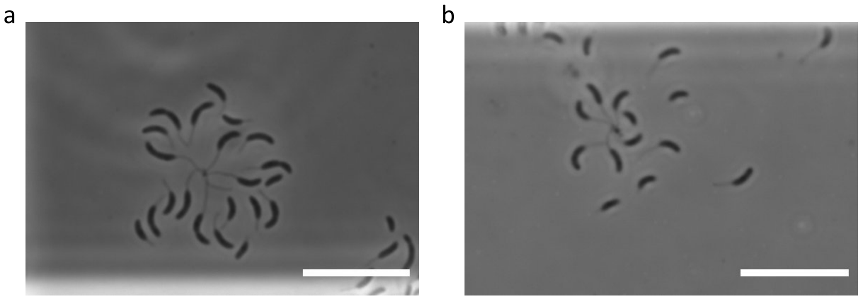


**Figure S1 | Rosette formation of flagella-lacking mutant on monomeric carbon sources.** Representative phase contrast microscopy images of Δ*flgH* strain growing on (**a**) xylose or (**b**) glucose for 16 hours. Scale bars correspond to 10 µm.

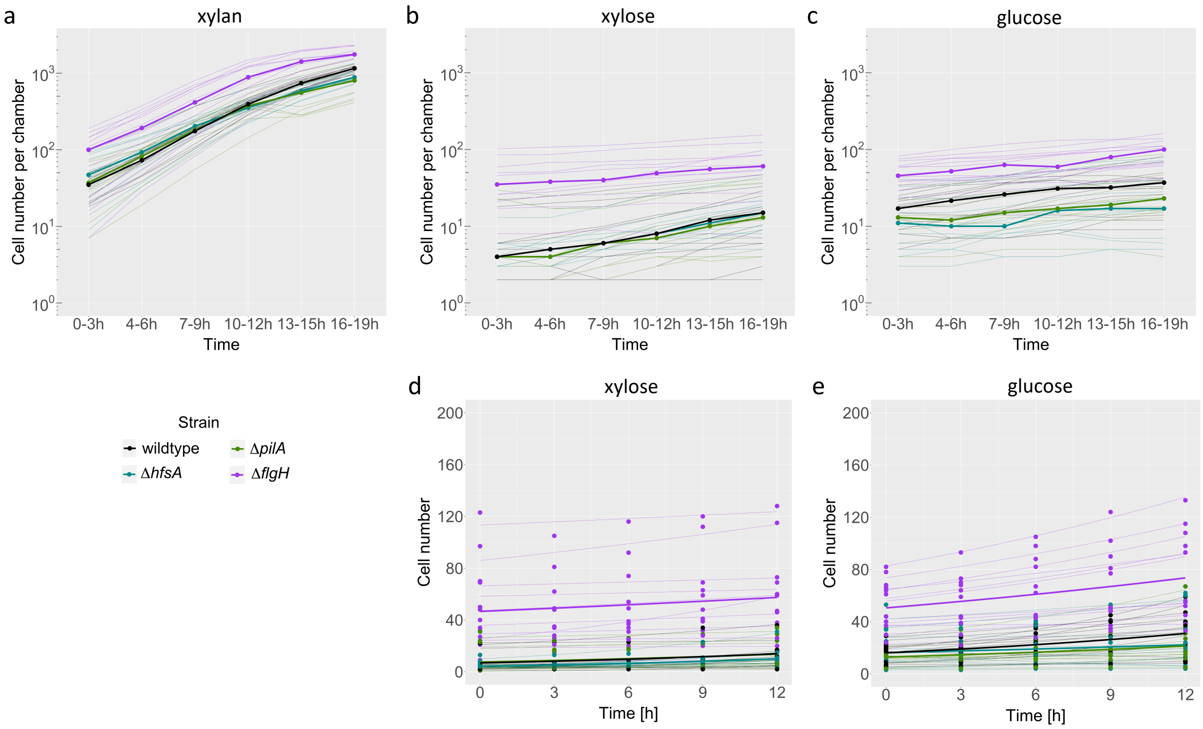


**Figure S2 | Cell number per chamber over time.** Median cell number per microfluidic growth chamber over time for (**a**) xylan, (**b,d**) xylose, and (**c,e**) glucose for each strain. Fine lines indicate the median cell number trajectories of single growth chambers. In **d** and **e**, the increase in cell number for each chamber was described with an exponential model

**(Supplementary Table S1)** and shown by a thin line (xylose: $R^2$ = 0.02-0.97, glucose: $R^2$ = 0.01-0.99). Thick lines depict the exponential regression lines using the mean parameter values determined in the model functions for the individual chambers. The graphs include data from 12 chambers, 4 per each of the 3 biological replicates, for each strain and carbon source.

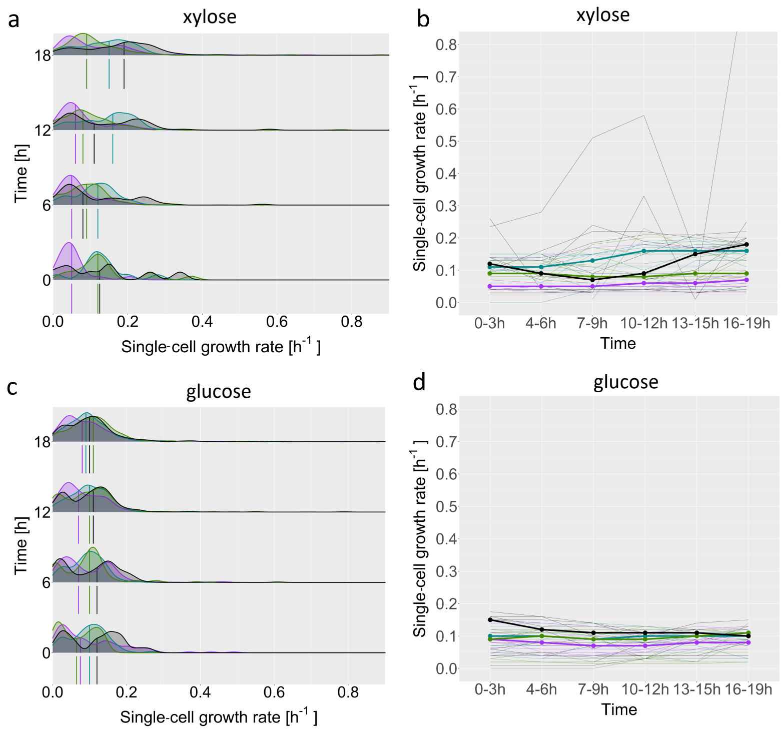


**Figure S3 | Single-cell growth rates on the monosaccharides xylose and glucose.** Single-cell growth rates over time on (**a,b**) xylose and (**c,d**) glucose for the different strains. Density plots show the distributions of single-cell growth rates at four timepoints, i.e. 0 h, 6 h, 12 h, and 18 h on (**a**) xylose and (**c**) glucose for the different strains. The vertical lines within and below the density functions depict the median values for each strain. Median values for single-celll growth rates over time for (**b**) xylose and (**d**) glucose for each strain (**Supplementary Table S3-5**). Thin lines indicate the median single-cell growth rate trajectories of individual chambers. All graphs include data from 12 chambers, 4 per each of the 3 biological replicates, for each strain and carbon source.

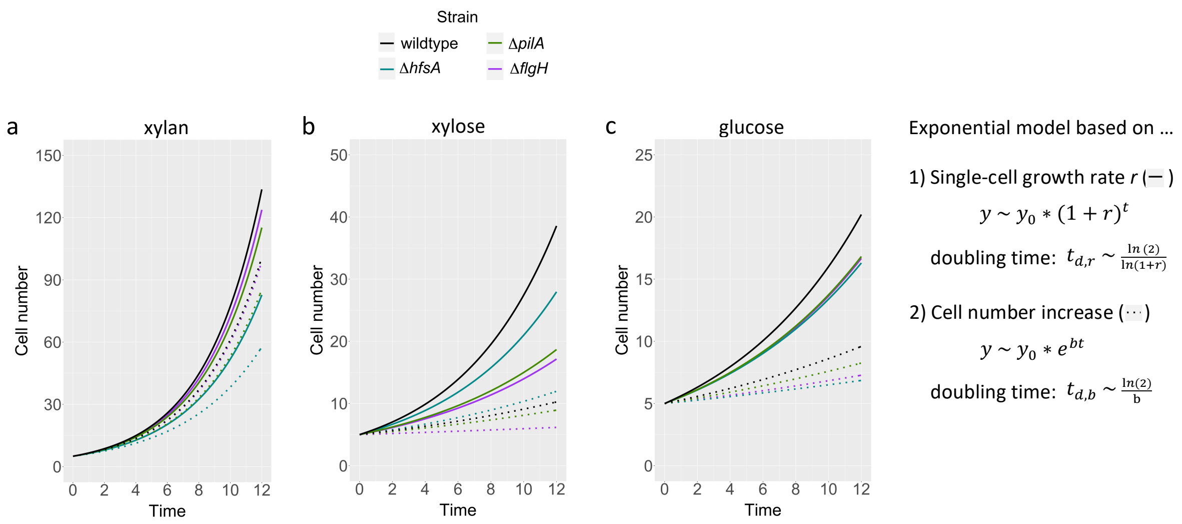


d

| | xylan | | | xylose | | | glucose | | |
|---|---|---|---|---|---|---|---|---|---|
| **Strain** | **r mean** | **b mean** | $\frac{t_{d,b}}{t_{d,r}}$ | **r mean** | **b mean** | $\frac{t_{d,b}}{t_{d,r}}$ | **r mean** | **b mean** | $\frac{t_{d,b}}{t_{d,r}}$ |
| wildtype | 0.315 | 0.250 | 1.09 | 0.186 | 0.060 | 2.85 | 0.123 | 0.054 | 2.14 |
| ΔpilA | 0.299 | 0.236 | 1.11 | 0.116 | 0.048 | 2.27 | 0.106 | 0.042 | 2.42 |
| ΔhfsA | 0.263 | 0.204 | 1.15 | 0.154 | 0.073 | 1.97 | 0.104 | 0.026 | 3.75 |
| ΔflgH | 0.307 | 0.249 | 1.08 | 0.108 | 0.017 | 5.92 | 0.106 | 0.031 | 3.21 |

**Figure S4 | Exponential growth models based on either single-cell growth rates or cell number increase per chamber.** Theoretical exponential increase of cell number per chamber over time *t* for (**a**) xylan, (**b**) xylose, and (**c**) glucose for each strain. A starting cell number $y_0$ of 5 cells was choosen for calculation. Regression lines are either based on the mean single-cell growth rate *r* of the first 12 hours (solid line) or the mean growth parameter *b* determined from the cell number counts over the first 12 hours (dotted line). The specific values for *b* and *r* as well as the doubling time ratios are summarized in **d**.

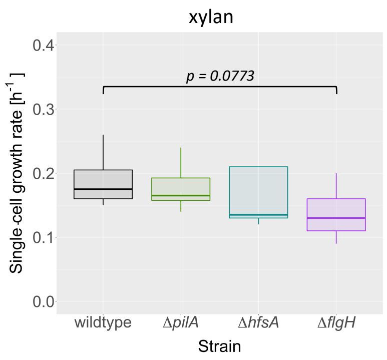


**Figure S5 | Comparison of single-cell growth rates between strains growing on xylan.** Boxplot showing the median, 25% quantile, and 75% quantile for cells growing on xylan (**b**). Statistical analysis **(Supplementary Table S6)** was performed using ANOVA (F = 2.2241, $p$ = 0.186)**.** For further pairwise comparison of the strains, we used the Tukey's multiple comparison test **(Supplementary Table S7)**. The data stems from 12 chambers, 4 per each of the 3 biological replicates, for each strain.

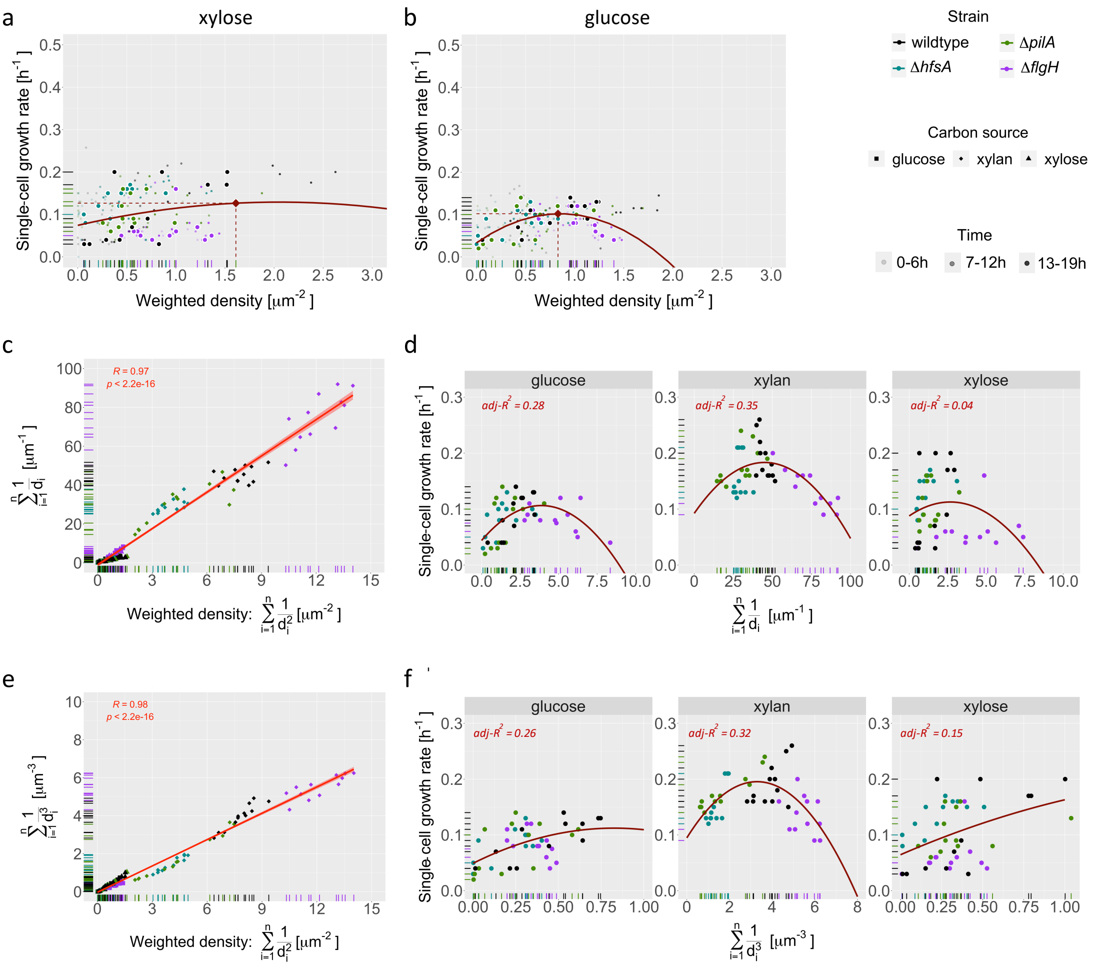


**Figure S6 | Testing robustness of the correlation between single-cell growth rate and cell density.** Median values of single-cell growth rate and weighted density plotted against each other for (**a**) xylose and (**b**) glucose. Larger points with white border represent time-averages of single growth chambers. Smaller points show single growth chambers at a specific time window indicated by the degree of transparency. The distribution of the median chamber values was fitted using a quadratic regression model (**Supplementary Table S8**) and depicted in red (xylose: adj-R2 = 0.02, glucose: adj-R2 = 0.31). The maxima of the quadratic regression line is indicated with a diamond shaped point in red (xylose: ymax = (1.61,0.13), glucose: ymax = (0.83,0.1)). Spearman's rank correlation between cell density measures using distance *d* (**c**) or *d* cubed (**e**) and the weighted density based on *d* squared. Each point represents the median value of a single growth chamber. Median values of single-cell growth rate plotted against the density measures based on *d* (**d**) or based on *d* cubed (**f**) for glucose, xylan, and xylose. The correlation between single-cell growth rate and the cell density measure was fitted using a quadratic regression model **(Supplementary Table S8)** and depicted in red. The data stems from 12 chambers, 4 per each of the 3 biological replicates, for each strain.

### SUPPLEMENTARY MOVIES

**Supplementary Movie M1 (separate file) |** Representative segmented and tracked time-lapse of *C. crescentus* CB15 wildtype growing on xylan. Images were captured every 7 minutes. Scale bar corresponds to 10 µm.

**Supplementary Movie M2 (separate file) |** Representative segmented and tracked time-lapse of *C. crescentus* CB15 ∆*pilA* growing on xylan. Images were captured every 7 minutes. Scale bar corresponds to 10 µm.

**Supplementary Movie M3 (separate file) |** Representative segmented and tracked time-lapse of *C. crescentus* CB15 ∆*hfsA* growing on xylan. Images were captured every 7 minutes. Scale bar corresponds to 10 µm.

**Supplementary Movie M4 (separate file) |** Representative segmented and tracked time-lapse of *C. crescentus* CB15 ∆*flgH* growing on xylan. Images were captured every 7 minutes. Scale bar corresponds to 10 µm.

**Supplementary Movie M5 (separate file) |** Representative segmented and tracked time-lapse of *C. crescentus* CB15 wildtype growing on xylose. Images were captured every 7 minutes. Scale bar corresponds to 10 µm.

**Supplementary Movie M6 (separate file) |** Representative segmented and tracked time-lapse of *C. crescentus* CB15 ∆*pilA* growing on xylose. Images were captured every 7 minutes. Scale bar corresponds to 10 µm.

**Supplementary Movie M7 (separate file) |** Representative segmented and tracked time-lapse of *C. crescentus* CB15 ∆*hfsA* growing on xylose. Images were captured every 7 minutes. Scale bar corresponds to 10 µm.

**Supplementary Movie M8 (separate file) |** Representative segmented and tracked time-lapse of *C. crescentus* CB15 △*flgH* growing on xylose. Images were captured every 7 minutes. Scale bar corresponds to 10 µm.

**Supplementary Movie M9 (separate file) |** Representative segmented and tracked time-lapse of *C. crescentus* CB15 wildtype growing on glucose. Images were captured every 7 minutes. Scale bar corresponds to 10 µm.

**Supplementary Movie M10 (separate file) |** Representative segmented and tracked time-lapse of *C. crescentus* CB15 △*pilA* growing on glucose. Images were captured every 7 minutes. Scale bar corresponds to 10 µm.

**Supplementary Movie M11 (separate file) |** Representative segmented and tracked time-lapse of *C. crescentus* CB15 △*hfsA* growing on glucose. Images were captured every 7 minutes. Scale bar corresponds to 10 µm.

**Supplementary Movie M12 (separate file) |** Representative segmented and tracked time-lapse of *C. crescentus* CB15 △*flgH* growing on glucose. Images were captured every 7 minutes. Scale bar corresponds to 10 µm.
